# Belgian endive-derived biostimulants promote shoot and root growth *in vitro*

**DOI:** 10.1101/2022.03.07.481290

**Authors:** Halimat Yewande Ogunsanya, Pierfrancesco Motti, Jing Li, Hoang Khai Trinh, Lin Xu, Nathalie Bernaert, Bart Van Droogenbroeck, Nino Murvanidze, Stefaan Werbrouck, Sven Mangelinckx, Aldana Ramirez, Danny Geelen

## Abstract

Recovering biostimulant compounds from by-products of crops is a promising strategy to add value, enhance sustainability, and increase the environmental safety of the agricultural production chain. Here, we report consistent root and shoot growth-stimulating bioactivity present in water-based extracts from Belgian endive forced roots (*Cichorium intybus* var. *foliosum*) over two consecutive harvest years. The shoot and the primary root of *in vitro* cultivated *Arabidopsis thaliana* treated with Belgian endive extract were about 30% increased in size compared to plants grown under control conditions. The ornamental species *Plectranthus esculentus* also showed enhanced *in vitro* shoot and root growth, suggesting bioactivity on a broad range of species. Fractionation of the Belgian endive extracts into aqueous and organic subfractions coupled with bio-activity measurements showed that the principal root and shoot growth-promoting ingredients are primarily water-soluble. NMR-based characterization of the bioactive aqueous fractions revealed the presence of predominantly sugars and organic acids. Malate and sugars were abundant and common to all water fractions, suggesting these molecules contributed to the growth stimulation phenotype. The findings indicate that Belgian endive roots are a source for the development of organic waste-derived biostimulants with potential for application in tissue culture and putatively for soil-grown crop production.

## 1. Introduction

Agriculture is under pressure because of climate change leading to more severe, more frequent, and longer periods of abiotic stress, causing substantial crop losses. At the same time, there is the desire to safeguard the natural environment by avoiding the use of synthetic agrochemicals. Researchers and industry are developing alternative methods and are looking for natural products termed biostimulants to improve crop resilience and yield ^1,2^. Most studies indicate that biostimulants enhance nitrogen metabolism ^3–5^ or improve nutrient and water uptake ^6^ or contain precursor molecules boosting metabolism. What most studies show, however, is that more pronounced effects are recorded under conditions of stress ^7–10^.

In general, biostimulants on the market are of organic origin and categorized as protein hydrolysates, polysaccharides, seaweed extracts, fulvic and humic acids, botanical extracts, and microorganisms ^11^. Many non-microbial biostimulants contain amino acids and organic acids. For instance, protein hydrolysates derived from plants are mixtures of amino acids and peptides that have shown positive effects on plant productivity and tolerance to abiotic stress ^2,5,12^. The majority of organic biostimulants are mixtures of different compounds and often combined with fertilizers, which makes it very difficult to assess whether the fertilizer, the biostimulant, or their combination are the cause for the trait or yield improvement ^13^. Apart from the added fertilizer, organic biostimulants are complex mixtures of biochemical compounds that separately may not exert the activity, raising the possibility that multiple compounds act together or in synergy to stimulate plant growth ^14^. Plant growth and performance are highly complex, and a combination of bioactive compounds may affect many processes.

Biostimulants are produced from various organic sources which are stable, cheap, and available in large quantities, preferably on a year-round basis. Agricultural and food processing by-products ^15^ constitute an alternative source of biostimulants, of increasing interest ^3,16,17^. In the view of sustainability and a circular economy, agricultural by-products are a valuable source to exploit for bioactive compounds and biostimulant production. For example, protein hydrolysates are derived from enzymatic, thermal, or chemical hydrolysis of proteins from by-products from the agriculture industries such as animal by-products, the biomass from tomato greens ^18^, rapeseed, apple seeds, and rice husk by-product ^19^. Other examples of organic wastes or by-products currently valorized as plant biostimulants are vermicompost ^7,20^ and compost tea ^21^. Also, aqueous extracts of by-products from fennel, lemon, brewer’s spent grain of barley, etc. have been investigated as sources of biostimulants ^17,22^.

*Cichorium intybus* var. *foliosum* is a popular crop in Europe covering about 95% of the world’s production with Belgium being the top producer in Europe ^23^, where it is cultivated as a vegetable crop grown for its etiolated leaves, known as Belgian endive (or witloof or chicon), and red endive (or radicchio), the latter of which is mostly cultivated in Italy. The leaves are produced from roots that are “forced” to sprout in the dark at 16–20 °C for about 21 days. The Belgian endive roots are harvested from the field and then stored in a cold room prior to forcing. Currently, the forced roots by-products are sometimes mixed with animal feed ^24^.

In this paper, we present a first report on the biostimulant activity of Belgian endive forced root by-product. We show that extracts prepared from the Belgian endive by-product (forced roots) promote root and shoot growth of *Arabidopsis thaliana* seedlings and *Plectranthus esculentus* explants cultivated *in-vitro.* Fractionation of the Belgian endive root extract revealed that the aqueous subfractions were enriched in both roots and shoot stimulating compounds. The aqueous fractions contained malate, multiple sugars choline, and primary metabolites that may enhance root and shoot growth in vitro. Forced Belgian endive roots, which is a stable, cheap, and abundantly available by-product in the center of Europe, have therefore potential to be developed into a biostimulant for promoting tissue cultured plants and thereby may contribute to the zero waste and circular economy concept.

## 2. Materials and Methods

### 2.1. Extraction and fractionation of biostimulants from Belgian endive forced roots

#### 2.1.1. Plant material pre-processing treatment

Belgian endive forced roots (*Cichorium intybus* var. *foliosum*) harvested in 2018 and 2020 were supplied by Versalof (Steenhuffel, Belgium) and further processed at the ILVO’s Food Pilot plant (Flanders Research Institute for Agriculture, Fisheries and Food, Melle, Belgium). First, forced roots were washed in cold water to remove the remaining soil. The outer ends of the roots were removed at the top and bottom. Further, the roots were julienned into 5 cm long and 2.5 × 2.5 mm width slices, using a Robot Coupe (CL50 Ultra, Mont-Sainte-Geneviève, France). The cut roots were placed in a hot air oven (60 °C, 6–8 h) to dry to a moisture content below 10%. Dried samples were milled by using a ring sieve size 0.5 mm (Ultra centrifugal mill ZM 200, RETSCH, Haan, Germany) to obtain a powder which is, in turn, used to produce water (HO), ethanol (EH), ethyl acetate (EA), and hexane (HE) extracts.

#### 2.1.2. Extraction procedure

Three and a half kilograms of dried forced roots powder were mixed with 31.8L of water. The resulting solid/water mixture (of approximately 35L) was incubated while stirring for 2h at 80°C in a so-called “Stephan’s apparatus”. Solid and liquid phases were separated by passing the mixture through 1000 μm and 100 μm vibrating sieves. The liquid phase obtained in this way constitutes the “liquid water extract (HO)”, which was concentrated to 13 L, aliquoted, and frozen at −20°C until further use. The resulting solid phase of the water extraction was then subjected to three sequential organic solvent extractions: ethanol (EH), ethyl acetate (EA), and hexane (HE) (Figure S1).

First, the solid left-over material from the first extraction was mixed and incubated with 51 L of ethanol at 60°C for 1 hour twice. After incubation, the solid/ethanol mixture was partitioned using a Buchner funnel. This procedure yielded the “liquid ethanol extract” and a new solid leftover phase. The ethanol contained in the “liquid ethanol extract” was evaporated *in vacuo* to dry ness and stored at −20°C as “dried ethanol extract (EH)”. The solid residue from the ethanol extraction went through an ethyl acetate (EA)extraction, and a subsequent hexane (HE) extraction, similarly as described before for the ethanol extraction. At the end of the whole extraction procedure, a liquid water extract (HO), and three solid organic solvent extracts (EH, EA, and HE) were obtained (Figure S1).

#### 2.1.3. Fractionation procedure

Prior to the fractionation, 11 L of 2018 HO extract previously obtained was concentrated to a final volume of 2 L, using a SpeedVac vacuum concentrator. The concentrated HO extract was divided into four 500 mL aliquots and the pH was adjusted to either pH 3 (aliquots 1 and 3) or pH 10 (aliquots 2 and 4) by using HCl or KOH, respectively. To proceed with the liquid-liquid fractionation, the volume of each aqueous aliquot was brought up to 1 L by the addition of water and later mixed with 2.2 L of either, ethyl acetate (aliquots 1 and 2) or toluene (aliquots 3 and 4) (Table S1). After mixing, and to speed up the separation process, organic and aqueous phases were separated by centrifugation (Eppendorf). The whole procedure yielded eight fractions, four organic fractions (F1: ethyl acetate-pH3; F2: ethyl acetate-pH10; F3: toluene-pH3; F4: toluene-pH10) and four aqueous fractions (F5: aqueous residue of F1; F6: aqueous residue of F2; F7: aqueous residue of F3; F8: aqueous residue of F4). The organic solvent (ethyl acetate (EtAc) and toluene (To)) contained in F1, F2, F3, and F4 was later removed by evaporation *in vacuo,* resulting in four organic/dried fractions. Aqueous fractions F5, F6, F7, and F8 remained intact and were frozen at −20°C until further use.

### 2.2. Biostimulants bioassays

#### 2.2.1. In-vitro Arabidopsis thaliana rooting and shooting bioassay

The four extracts (HO, EH, EA, HE) and the eight fractions (F1 – F8) prepared as described above (Items 2.1.2 and 2.1.3) were incorporated in MS (Murashige and Skoog) basal medium [1.5 g/L MS basal salt (Duchefa), 5 g/L sucrose, 0.5 g/L MES monohydrate, 8.0 g/L plant tissue culture agar (Duchefa); pH 5.7] at different concentrations. The HO extract and aqueous fractions F5 to F8 were diluted to the concentrations shown in Table 1. The solid extracts (EH, EA, and HE) and organic fractions F1 to F4 were dissolved in pure dimethylformamide (DMF) and then diluted 10-, 100-, and 1000-times using growth medium (Table 1), with the highest concentration containing 0.05% DMF. Water and DMF (0.05%) were used as controls for aqueous extract/fractions and organic extract/fractions, respectively. *Arabidopsis thaliana* Col-0 seeds (collected in HortiCell lab) were sterilized in 1.5 ml Eppendorf tubes using vapor-phase seed sterilization. Briefly, the seed-containing tubes were placed in a rack and the rack was placed in a desiccator in a fume hood. Next to the rack is a beaker containing 100 mL of bleach, to which 3 mL of hydrochloric acid was added. The seeds were exposed for 4 hours, after which they were transferred to a sterile laminar flow hood and left open for 2 h. Sterile *Arabidopsis* seeds were germinated and etiolated as previously described ^25^. Etiolated seedlings were transferred to freshly prepared, treatment (MS medium with extracts or fractions) and control (MS medium without extracts/fractions) square plates (12 × 12 cm). Root morphology (primary root length) and shoot morphology (leaf area) traits were examined and recorded by digital photography after 10 days of incubation (warm white light, 70 μmol.m^-2^.s^-1^ intensity, 16 h light/8 h dark photoperiod at 21 °C). Using the Fiji software ^26^, images were used for the scoring of primary root length as described by ^25^ and the leaf area. An Olympus binocular microscope was used for the lateral root and adventitious root numbers manual counting.

**Table 1.**
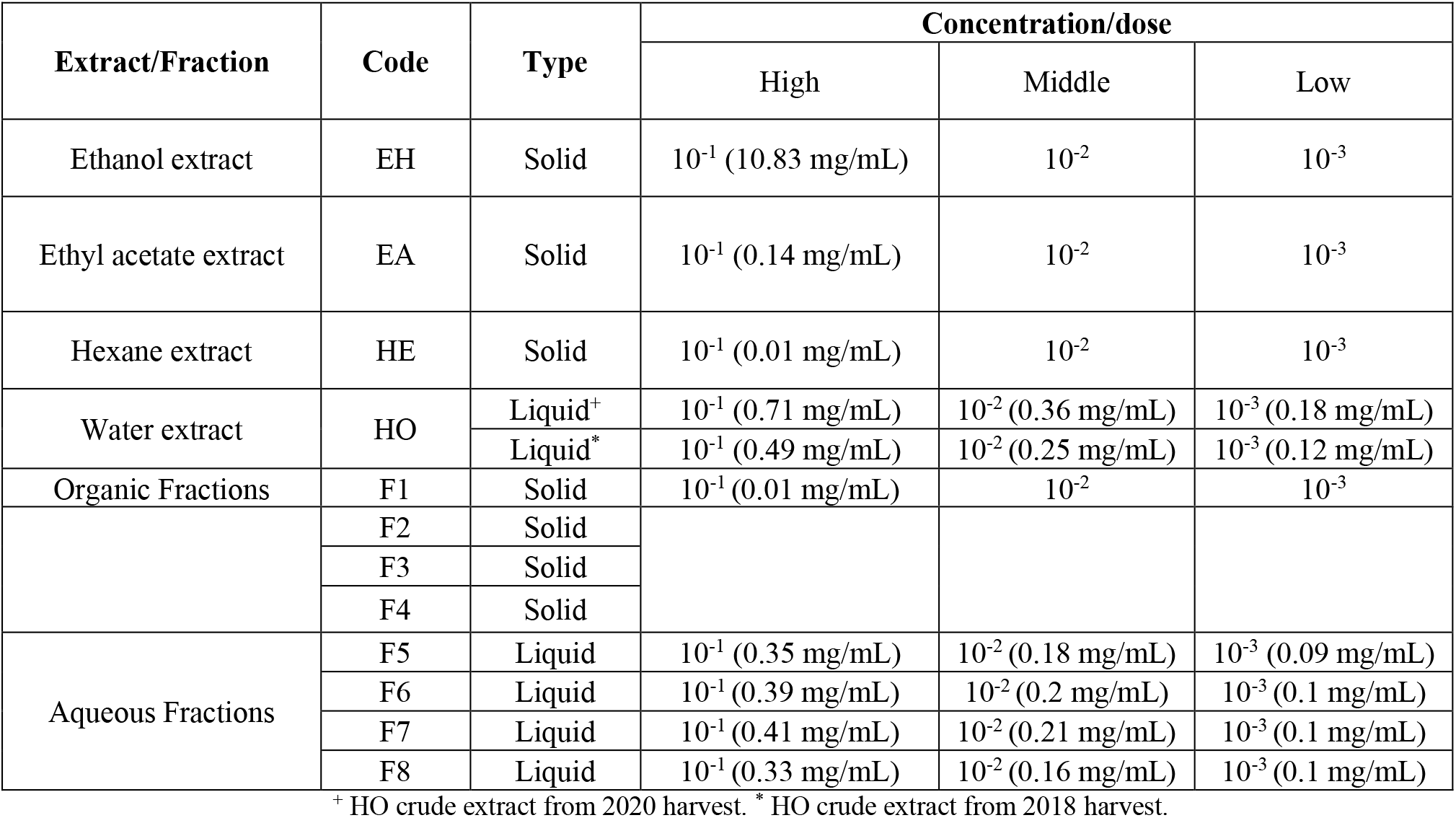
Extract and fraction concentrations used in the *Arabidopsis thaliana* root and shoot bioassays.

#### 2.2.2. In-vitro *Plectranthus esculentus* rooting and shooting bioassay

Glass jars of 350 ml were filled with 100 mL Murashige & Skoog medium with a half concentration of NH_4_NO_3_ and KNO_3_, including microelements and vitamins. This basal medium was supplemented with 30 g/L sucrose, 7 g/L agar-agar, and either the Belgian endive forced root HO extract, or its derived fractions (F1 - F8). The HO liquid extract was tested at two dilutions, 1/100 and 1/1000 (10^-2^ and 10^-3^), the organic fractions (F1 - F4), which were solid, were tested at a concentration of 10mg/L (10^-5^), and the liquid aqueous fractions (F5 - F8) were tested at a dilution of 1/100 (10^-2^). Controls consisted of no additions of extract or fractions. Plants were cut into uniform 1 cm length including leaves and two axillary buds, opposite of each other, and 10 explants per jar (two jars per treatment = 20 explants) were cultured. Cultures were maintained under cool fluorescent light, provided by PHILIPS master TLD 36 W 830 Reflex ECO (40 μmol m^-2^ s^-1^ PAR) 16 h light and 8 h dark photoperiod at 22 ±2 °C. After 3 weeks the presence of new shoots, roots, and the length of the root, was assessed.

The plant materials used in these experiments are in accordance with the rules of the national and international guidelines and legislation.

### 2.3. NMR sample preparation, acquisition, and processing

The aqueous fractions F5-F8 of the HO extract of 2018 and 2020 HO crude extract were dried *in vacuo* using the rotavapor and high vacuum apparatus. Of each dried sample, 20 mg was dissolved in 450 μL of D_2_O buffered with KH_2_PO_4_ (90 mM, pH 7, Sigma Aldrich) and 100 μL of a 5mM DSS solution in D_2_O. D_2_O and DSS provided a field frequency lock and chemical shift reference (^1^H δ 0.00 ppm) respectively. NMR experiments were performed using a Bruker AVANCE III spectrometer, equipped with 1H/BB z-gradient probe (BBO, 5 mm) for the ^1^H. ^13^C, ^1^H-^1^H COSY, and ^1^H-^1^H phase-sensitive TOCSY experiments.

The ^1^H and ^13^C NMR spectra were measured at 400 and 100.6 MHz, respectively. All spectra were acquired through the standard pulse sequences available in the Bruker pulse program library. Spectral data were all processed with Bruker TopSpin version 4.1.3. Exponential window multiplication of the FID, Fourier transformation, and phase correction were performed using Bruker AU programs proc_1d for 1D experiments and proc_2dsym for 2D experiments. Identification of the compounds was performed manually by comparing the ^1^H NMR spectra with the spectra available in the HMDB library, with the help of Chenomx NMR suite 9.0 software (Chenomx Inc., Canada). 2D NMR spectra were calibrated and visualized with Bruker TopSpin 4.1.3 to confirm the ^1^H NMR assignments. ^13^C NMR data were visualized and analyzed with ACD/Spectrus Processor software (ACD/Labs, Canada), and the assignment was performed by comparison with HMDB ^27^ and BMRB ^28^ databases.

The ^1^H NMR data quantification was attained by acquiring the integration values of selected resonance signals of all the identified metabolites with the reference integration value of the resonance signal at 0.00 ppm of DSS set to 9. This was manually done using Bruker TopSpin version 4.1.3. Normalization for each metabolite was done by dividing the corresponding integration value of the resonance signal of each metabolite by its corresponding number of protons. The relative abundance of the metabolites was then determined by dividing the corresponding normalized integration value of each metabolite by the total of the normalized integration values of all identified metabolites in the respective fractions. These relative abundances were represented as percentages. All calculations were done using Excel (Microsoft Inc.).

### 2.4. Statistical analysis

The data collected were plotted and analyzed using GraphPad Prism 8.0.2 and the results were expressed as average ± standard mean error (SEM). The differences between the mean values of treatments were analyzed using one-way ANOVA and two-way ANOVA followed by Dunnett’s and Tukey’s multiple comparison tests, respectively. A value of p < 0.05 was considered statistically significant. Principal component analysis (PCA) was performed using R 3.6.1 (R Foundation for Statistical Computing, Vienna, Austria).

## 3. Results

### 3.1. Biostimulant activity of crude extracts

#### 3.1.1. Effect of HO, EH, EA, and HE on *A. thaliana* root architecture

The *Arabidopsis* root system is composed of a long primary root from which lateral roots branch off. Additional roots that emerge from the hypocotyl (adventitious roots) are induced when the seedling is first cultivated in the dark and then transferred to the light. Biostimulant activity was determined by recording primary root length (PRL), the number of lateral roots (LR), and the number of adventitious roots (AR). A water extract of dried endive roots (HO) from the 2020 harvest was tested and found to significantly increase the primary root length (Figure 1a). The PRL was about 30% longer in the presence of low and mid doses of HO. A positive effect of HO was also observed on lateral root numbers at the same dose that increases the PRL (low and mid, Figure 1b). Longer primary roots have the propensity to form more lateral roots and we, therefore, calculated the LR index reflecting the LR density (number of LR per PRL). The LR density was consistent for all incubation conditions (Figure 1d) leading to the conclusion that we did not find evidence for endive extract stimulating LR induction. Likewise, HO treatment did not stimulate adventitious root formation and on the contrary, the AR number was lower upon low dose of HO treatment (Figure 1c). Root growth stimulation by HO was therefore most noticeable for the primary root (Figure S3).

**Figure 1.**
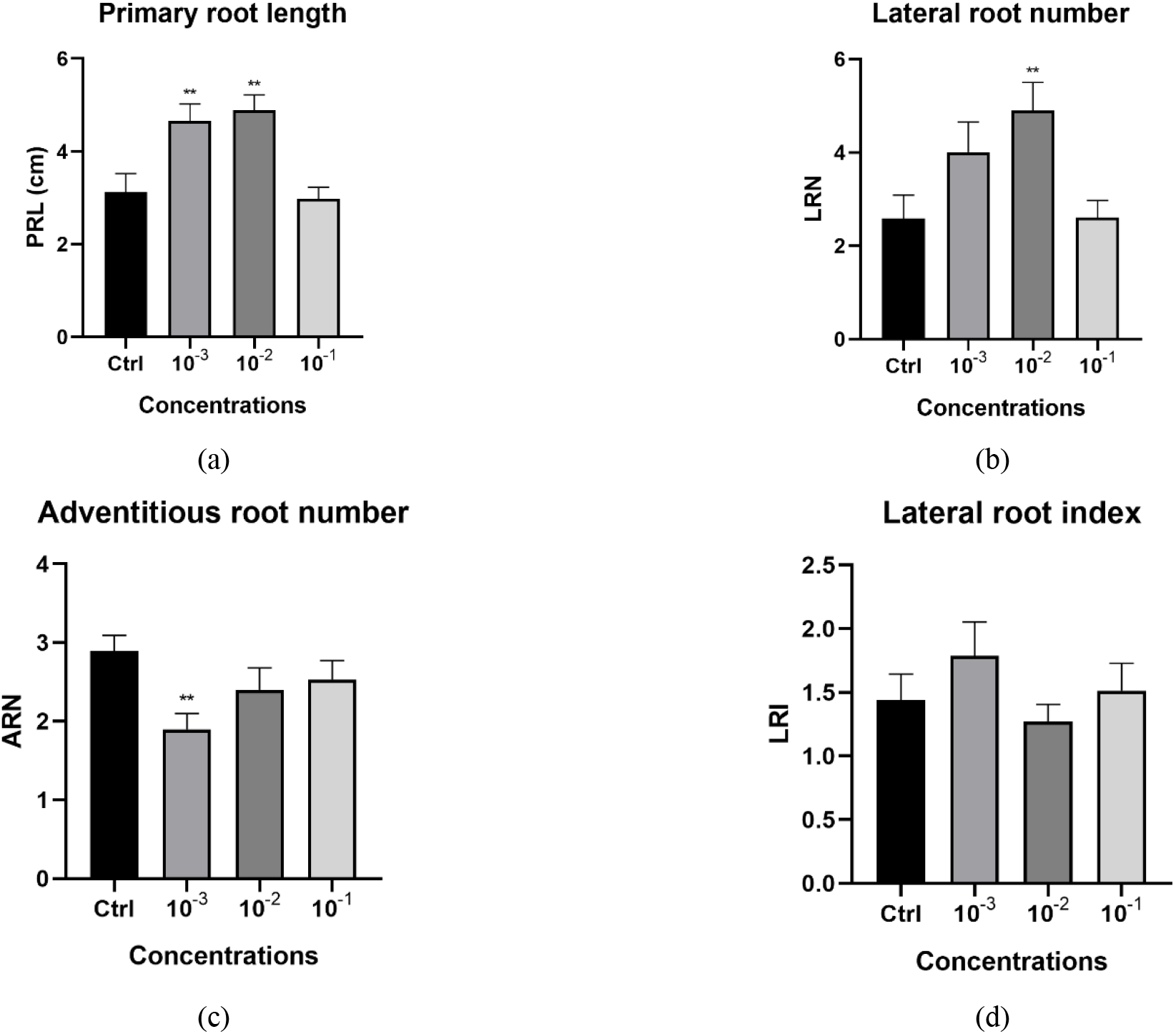
*Arabidopsis* root architecture stimulation upon treatment with low, mid, and high doses of HO. Graphical representations showing the effect on primary root length (a); on lateral root number (b); on adventitious roots numbers (c); and on the lateral root index (d). Data represent the average of three biological and ten technical replicates per bar (30 seedlings in total, 10 per replicate). Error bars represent standard mean error (SEM). Asterisks indicate significant differences between control and treatment according to Dunnett’s multiple comparison test (* p<0.05, ** p<0.01, *** p<0.001, **** p<0.0001). PRL: primary root length, LRN: lateral root number, ARN: adventitious root number, and LRI: lateral root index

Ethanol extract EH strongly inhibited primary root growth at the highest concentration, which was accompanied by a strong increase in the number of AR and a strong decrease in LR formed (Figure S2). EA and HE extracts showed in general more mild effects with a tendency to stimulate primary root growth (Figure S2a). EA at the highest concentration significantly reduced the number of LR (Figure S2b).

#### 2.1.2. Effect of HO, EH, EA, and HE on *A. thaliana* shoot growth

The shoot of *in-vitro* grown seedlings was substantially larger and greener when treated with HO (Figure 2a). Shoot growth was therefore quantified by analyzing the projected leaf area. Shoots of HO-treated plants were significantly larger than in the control plants with the mid dose treatment having the strongest effect of more than a two-fold increase in shoot area (Figure 2b). Although the increase in shoot area might be associated with the increase in PRL, this link was not observed in plants treated with organic extracts. Here, shoots growth was reduced by EH and EA treatments at the highest concentration despite the limited effect on PRL (Figures S2a & 2b)

**Figure 2.**
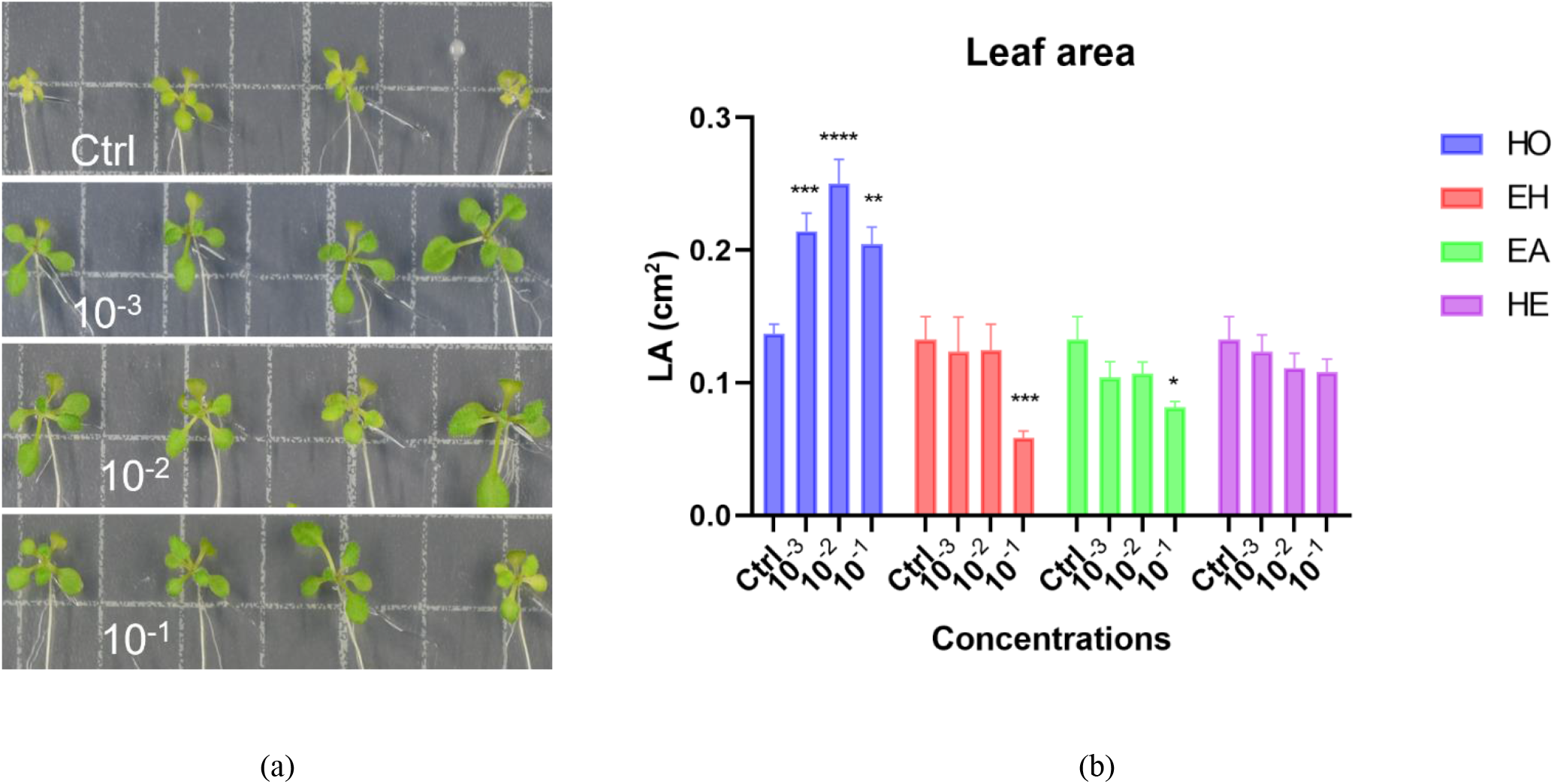
*Arabidopsis* shoot stimulation upon treatment with low, mid, and high doses of HO, EA, EH, and HE. (a) Images showing the leaf area of control (Ctrl), low, mid, and high dose of HO treated plants. (b) graphical representation of the effect of HO, EA, EH, and HE on the leaf area. LA: leaf area. Data represent the average of three biological and ten technical replicates per bar (30 seedlings in total, 10 per replicate). Error bars represent standard mean error (SEM). Asterisks indicate significant differences between control and treatment according to Dunnett’s multiple comparison test. (* p<0.05, ** p<0.01, *** p<0.001, **** p<0.0001).

### 3.2. Enrichment of endive forced roots bioactive ingredients by fractionation

#### 3.2.1. Effect of different fractions on the *A. thaliana* root architecture

A positive effect was obtained with diluted extract while the extract used at higher concentration was not active, suggesting that the growth stimulation is caused by growth-regulating compounds rather than e.g., the mineral content of endive extract. From the results of the crude extracts experiments conducted, only the HO extract significantly showed biostimulant activities by positively influencing the plant phenotypes examined. Therefore, a liquid-liquid fractionation of the HO extract was conducted to reduce the complexity of the extract. This fractionation yielded eight fractions, four organic fractions; F1 – F4, and four aqueous fractions; F5 – F8 (see Materials and Methods). These fractions were further subjected to bioactivity testing to determine which fractions retained the biostimulant activity of the original HO extract.

The eight fractions obtained from the fractionation of the endive water “HO” extract were subjected to bioactivity testing. The effect of the organic fractions (F1 – F4) on the root parameters varied. A significant effect of the organic fractions on the primary root length was observed only on plants treated with fractions F2 and F4 and at concentrations 10^-3^ and 10^-2^ dilutions (Figure 3a). Only fraction 4 significantly increased the lateral root number amongst the organic fractions, and this effect was observed at the lowest concentration tested (10^-3^ dilution, Figure 3c). All organic fractions except fraction 2 showed a significant increase in the number of adventitious roots. The significance was observed at the highest concentration tested (10^-1^ dilution, Figure 3e).

**Figure 3.**
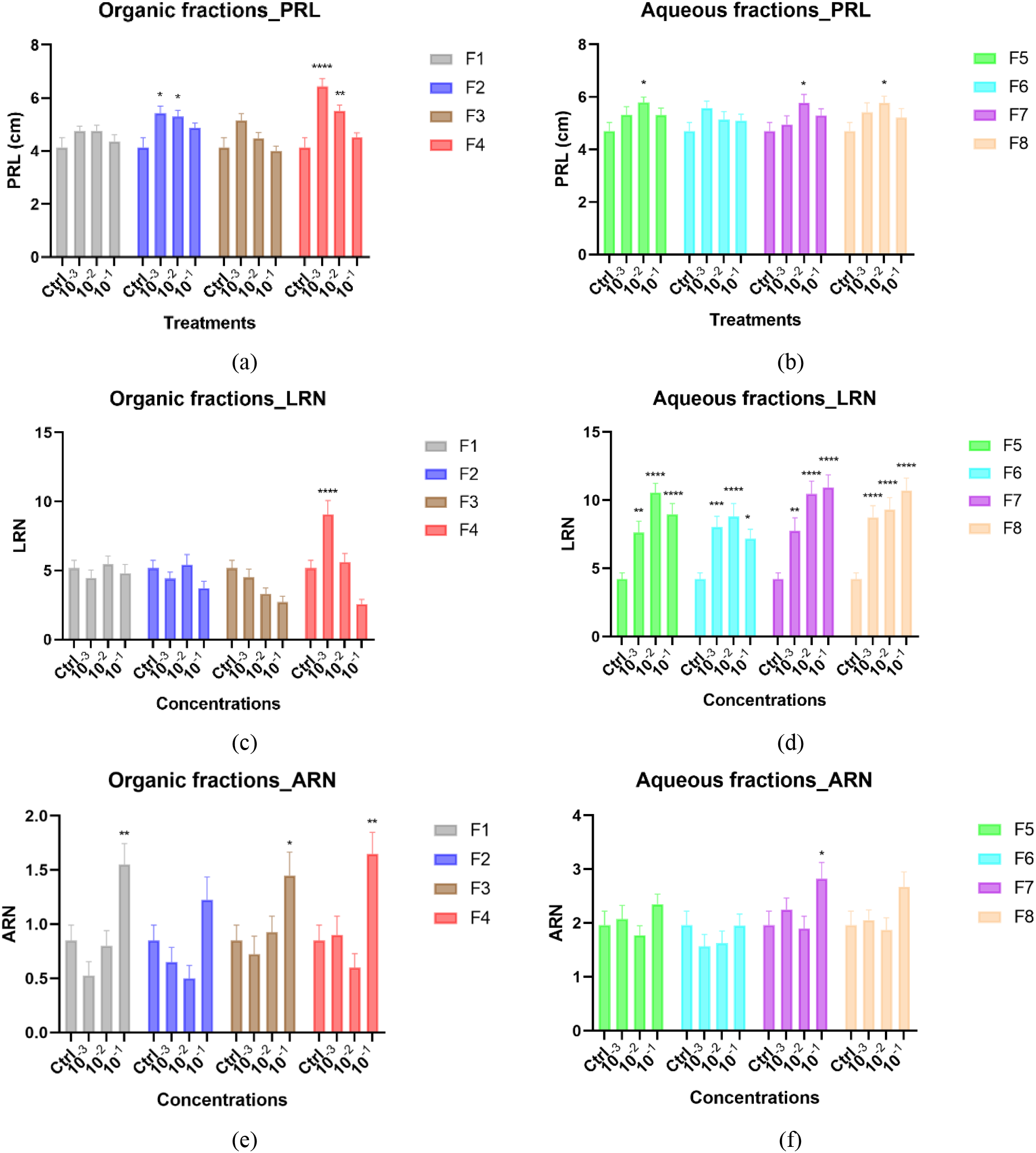
*Arabidopsis* root architecture stimulation upon treatment with different concentrations of the fractions from HO. Graphical representations showing the effects of the fractions on primary root length (a, b); on lateral root number (c, d); on adventitious roots numbers (e, f). Data represent the average of three biological and ten technical replicates per bar (30 seedlings in total, 10 per replicate). Error bars represent standard mean error (SEM). Asterisks indicate significant differences (* p<0.05, ** p<0.01, *** p<0.001, **** p<0.0001) between negative control (Ctrl) and treatments (doses) in respective groups (fractions) according to Tukey’s multiple comparison test. PRL: primary root length, ARN: adventitious root number, LRN: lateral root number, F1 – F8: fractions 1 to 8 denoted by different colors.

The aqueous fractions (F5 – F8) also showed varied effects amongst the root phenotypes examined, especially between the lateral roots and the adventitious roots. The length of the primary root was significantly increased by all aqueous fractions at mid dose except fraction F6 (Figure 3b). Unlike the effect of the organic fractions on the lateral root, all the aqueous fractions highly significantly increased the lateral root number with over a 100% increase (Figure 3d). Like the lateral root, an opposite effect was seen between the organic and aqueous fractions on the adventitious roots. Only aqueous fraction F7 increased the adventitious root number at high concentration (Figure 3f).

The fractions F4 and F7 positively influenced the three root phenotypes examined. The organic fractions maximally influenced the root length and lateral branching at 10^-3^ dilution, while aqueous fractions maximally influenced these root phenotypes at mid concentration. Adventitious root branching showed maximum effect at the highest concentration of three organic fractions and one aqueous fraction (Figure 3e,f).

#### 3.2.2. Effect of different fractions on the *A. thaliana* shoot growth

The leaf area was taken as a proxy to measure the impact of endive extracts and fractions on shoot growth. The strongest shoot growth promotion occurred when seedlings were treated with the aqueous fractions, increasing with the concentration applied (Figure 4b). The organic fractions also stimulated shoot growth, but here the optimal effect occurred at the intermediate dilution 10^-2^ for F1 and F2 and at the lowest dilution 10^-3^ for F3 and F4 (Figure 4a). Overall, the shoot growth-promoting compounds were hence not separated by the fractionation method applied.

**Figure 4.**
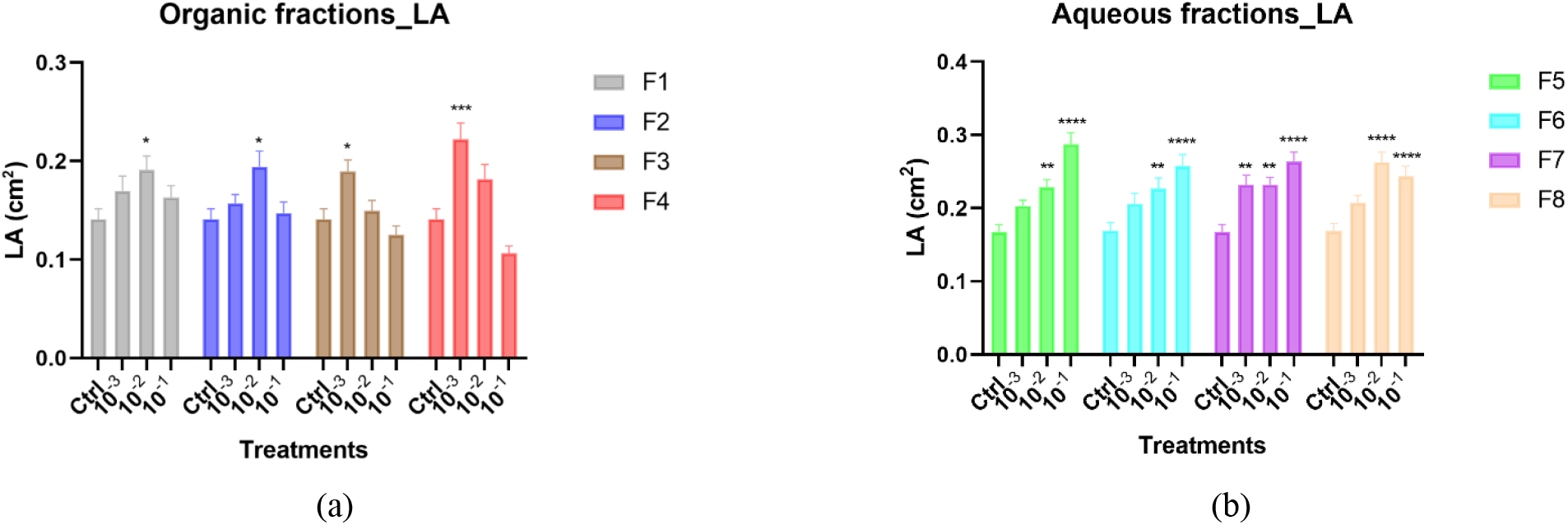
*Arabidopsis* shoot architecture stimulation upon treatment with different concentrations of fractions from HO. Graphical representations showing the effects of the organic fractions (a) and aqueous fractions (b) on leaf area. Data represent the average of three biological and ten technical replicates per bar (30 seedlings in total, 10 per replicate). Error bars represent standard mean error (SEM). Asterisks indicate significant differences (* p<0.05, ** p<0.01, *** p<0.001, **** p<0.0001) between negative control (Ctrl) and treatments (concentrations) in respective groups (fractions) according to Tukey’s multiple comparison test. LA: leaf area, F1 – F8: fractions 1 to 8 denoted by different colors.

Principal component analysis was conducted for the fractions data to see which phenotypes are more linked than others. The first two components of the PCA explained 71.9% of the variance in the dataset (PC 1: 42.6%; PC 2: 29.3%) (Figure S6). PC1 explained LRN (67.6%), PRL (69.3%) and partially LA (15.5%), while PC2 explained ARN (69.6%), LA (69.5%) and partially LRN (13.9%). From the analysis, PRL was positively correlated to LRN (64.6%) and LA (5.8%) and was negatively correlated to ARN (Figure S6). On the other hand, the shoot growth (LA) was positively correlated to all phenotypes (PRL: 5.8%, LRN: 14.6%, and ARN: 15.1%). This means that PRL and LR numbers are more highly correlated to each other than other phenotypes and LA is more correlated to AR numbers. Regardless of the correlations of the phenotypes, the plants treated with aqueous fractions showed enhanced root and shoot phenotypes than the plants treated with organic fractions.

### 3.3. Consistency of root and shoot growth-promoting activity over different endive harvest years

A recurrent problem with bioactivity analysis of products from natural resources is its reproducibility over separate extract preparations ^13^. Therefore, we compared the bioactivity of a water extract from endive roots harvested in 2018 with that of 2020 extracts which were used to generate data shown in Figure 1. The treatment with the 2020 endive extract resulted in a significant increase in PRL at low and mid doses, results that are comparable with those obtained with the 2018 extract (Figure S4a). The LR number was higher and the AR number lower in the experiments performed in 2018 compared with the 2020 experiments. Over this period, the shelf cooling system in the growth room was refurbished which we presume had an impact on the root development. Despite this inadvertent difference in impact on root branching, the same trend in LR increase and AR equivalency or reduction was recorded for both the 2018 and 2020 extracts (Figure S4b, c).

### 3.4. Root and shoot growth-promoting activity in Plectranthus esculentus

In view of the development of a growth-promoting biostimulant, we tested the impact of HO and fractions from 2018 on a second plant, *Plectranthus esculentus*, an ornamental plant species. To this end, we incubated *in-vitro* grown *P. esculentus* explants in the presence of different concentrations of HO. Figure 5a shows that compared to the water control, application of HO significantly stimulated the development of adventitious roots on *P. esculentus* shoot explants at 10^-3^ and 10^-2^ concentrations. Because shoot explants do not have a primary root, the PRL could not be determined, and instead, the length of the adventitious roots was analyzed. A significant increase in root length was observed at the lowest concentration applied (Figure 5b). As shoot formation is an important aspect of micropropagation, the formation of new shoots (Figure 5e) was also analyzed. HO treatment at the lowest concentration induced a higher number of shoots (Figure 5c), although not significantly different from the water control. Conversely, HO treatment at both concentrations significantly increased the length of the shoot internodes (Figure 5d).

**Figure 5.**
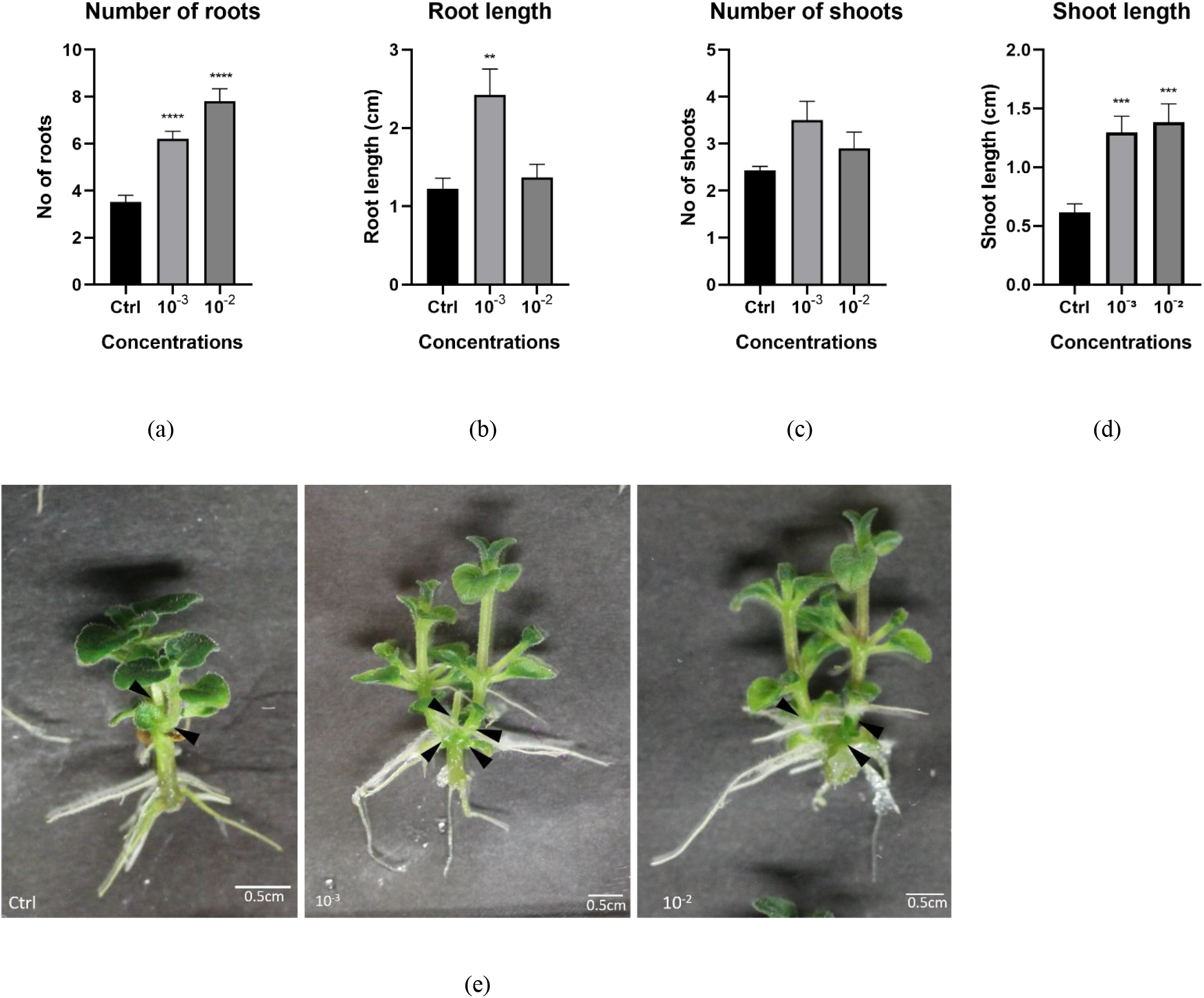
*P. esculentus* root and shoot architecture stimulation upon treatment with water (control) or with two concentrations (10^-3^ and 10^-2^) of HO. Graphical representations showing the effect of HO on (a) root numbers, (b) root length, (c) shoot numbers (black arrowheads), and (d) shoot length. (e) Images of the mature explants at the concentrations tested including control. Data represent the average of two biological and ten technical replicates per bar. Error bars represent standard mean error (SEM). Asterisks indicate significant differences between control and treatment according to Dunnett’s multiple comparison test (* p<0.05, ** p<0.01, *** p<0.001, **** p<0.0001).

The HO fractions were also tested for their growth-promoting effects on *P. esculentus* explants. The results showed that only the organic fraction F2 significantly stimulated the adventitious roots more than the water control (Figure S5a). On the other hand, none of the fractions significantly improved the formation of new shoots (Figure S5b).

### 3.5. NMR analysis

Manual inspection of the ^1^H NMR spectral data of fractions F5-F8 identified water-soluble metabolites including amino acids, organic acids, and sugars (Table 2). Comparison of the ^1^H NMR profile at 1-3.3 ppm with established databases (HMDB and BMRB) identified L-alanine, malate, and choline in all water fractions (F5-8). L-aspartate, citrate, formate and acetate were identified in F6 and F8. In addition, F6 contained lactate and F5 and F7 contained L-arginine, and F5, F6, and F7 contained fumarate. 2D NMR analysis (COSY and TOCSY), was instrumental in assigning signals at the 3.2-4.3 ppm range (Table 2). The overlap of resonance signals at this region prevented the identification of most peaks, yet the anomeric protons of sucrose, α-D-glucose, and β-D-glucose were identified (Table 2, Figure S7). The molecules identified in the ^1^H NMR mode were affirmed by ^13^C NMR analysis. The ^13^C NMR spectra in the range 62-110 ppm (Figure S8) showed that all fractions (F5-F8) generated similar resonance signals, suggesting a similar carbohydrate composition. Sucrose, α-D-fructose, and β-D-fructose signals were detected in the ^13^C NMR spectra; however, α-D-glucose and β-D-glucose were not detected (Table S2). Analysis of the ^13^C NMR spectra further identified L-arginine in fractions F5 and F7 and acetate in fractions F6 and F8. Based on previously reported spectral data 1-kestose was identified in all fractions (Table 2) ^29^.

The relative abundance (%) of the identified metabolites was determined by quantifying the ^1^H NMR data as described in section 2.3. The order in relative abundance was sugars > organic acids > amino acids > other compounds (Table 2). Comparatively, sugars were highly abundant in all the fractions, with α-D-glucose the least abundant sugar, except for F5, where its abundance was slightly higher than β-D-glucose (6.3 % and 4.9%). For the organic acids, malate prevailed in all fractions at varying abundance. Notably, in fractions F5 and F7, malate was the only organic acid present with relative abundances of 19% and 19.7%, respectively. In contrast, lactate was identified only in fractions F6 (18.8%) and F8 (1.1%). These findings align with malate fractionated in water during acidic (pH 3) extraction and lactate by basic (pH 10) extraction. Although detected in all the fractions, choline was consistently one of the least abundant metabolites.

To corroborate the finding that the bioactivity of the HO-treatment was consistent over different harvest years, ^1^H NMR analysis was conducted on 2020-HO crude extract. The relative abundance of the identified compounds in the 2018 aqueous fractions was reminiscent to that of the 2020 crude extract (Table 2). From the analysis, F5, F7 and 2020-HO crude all have about 70% saccharides, while F8 contains about 80% saccharides and F6 about 50%. The 2020-HO crude contained 19.4% malate (similar abundancy as F5 and F7) and the other organic acids, citrate (13.0%), lactate (4.2%), acetate (11.7%), fumarate (0.9%) and formate (0.4%) resembled that of F6 and F8. The choline abundancy in 2020-HO crude (1.7%) was higher as compared to the 2018-fractions (0.4-0.6%).

**Table 2.**
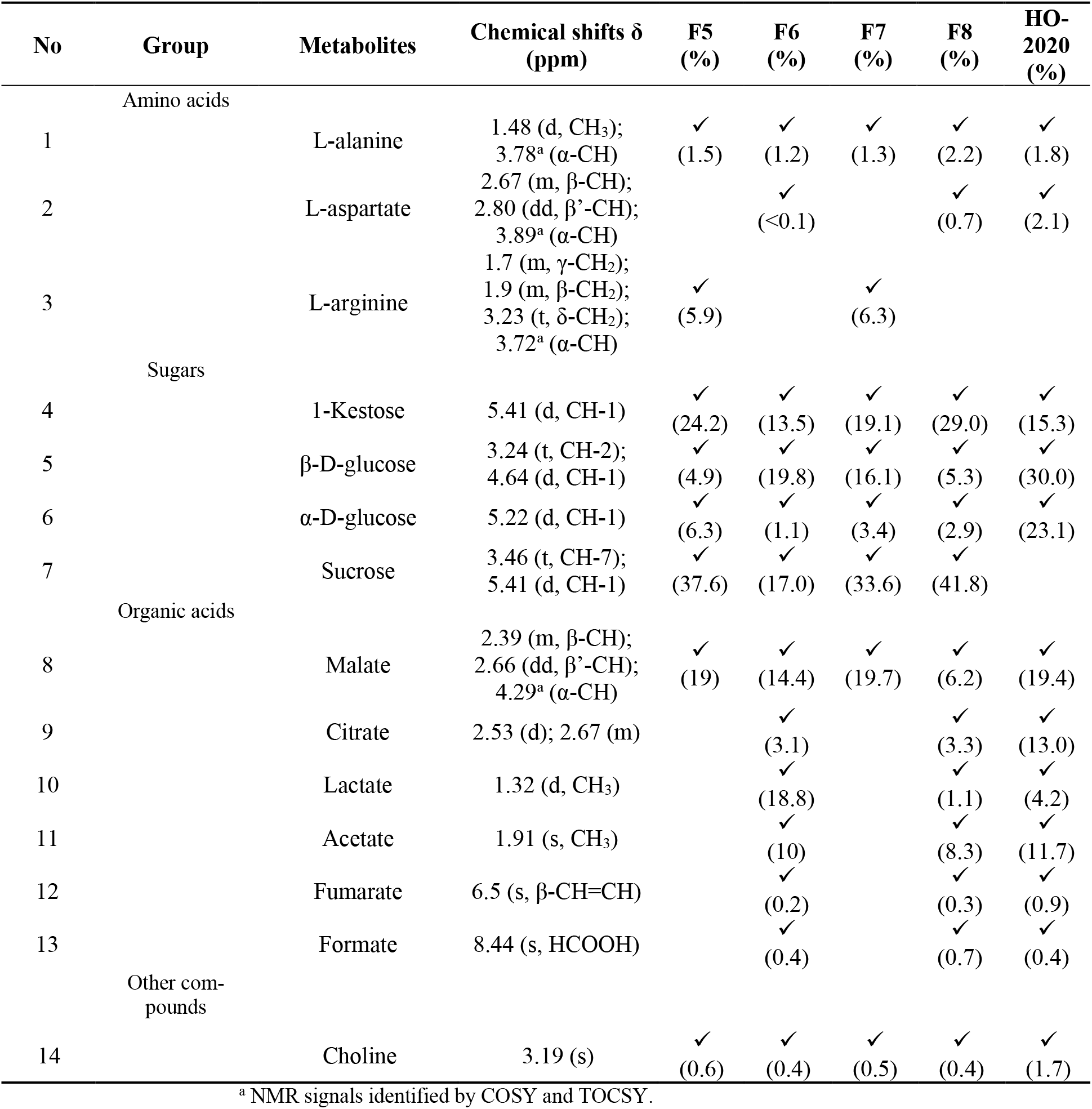
^1^H NMR (D_2_O, ref: DSS, pH 7, 400 MHz) chemical shifts of the identified metabolites, their presence, and relative abundances (%) in respective fractions F5-F8 and crude HO-2020.

## 4. Discussion

In this paper, we report the biostimulant activity of extracts produced from forced Belgian endive roots, a by-product from witloof production. Bioactivity consisted of the stimulation of shoot and root growth, which was recorded in two unrelated dicotyledonous species, suggesting a conserved mode of action. Plant biostimulants are typically mixtures of compounds that display bioactivity in a synergistic manner ^30^. The Belgian endive extracts studied in this work were also complex mixtures for which we anticipate additive, synergistic, and antagonistic interactions. Different sub-fractions showed growth stimulation of the shoot and root, indicating that multiple bioactive compounds are present in the Belgian endive extract. Since bioactivity was recorded in aseptic environments, the growth stimulation was direct and not associated with exogenous microbial interactions or biochemical conversions ^5,14,31^.

Aqueous fractions showed the strongest shoot growth effect at the highest concentration and the organic fractions showed an optimal effect at the intermediate concentration. The unsuccessful attempt to enrich for shoot growth-promoting compounds by two-phase solvent partitioning is remarkable. The possibility that multiple hydrophobic and hydrophilic bioactive compounds are present in the HO extract cannot be excluded, however, in view of the consistent dose response effects with both aqueous and organic fractions, we speculate that solvent partitioning based on hydrophobicity and the pH-dependent ionic character was not adequate for the complete separation of bioactive compounds. Specific physicochemical properties of the bioactive compound(s) may explain the lack of separation. Indeed, complex amphipathic molecules may adopt different structural configurations that prevent polar partitioning ^32^. Alternative fractionation strategies may address this issue in future experiments. A general activity of biostimulants is the improvement of photosynthesis that results in higher carbohydrate accumulation and increased biomass formation ^33–36^. The photosynthetic capacity of the plants was not measured in our *in vitro* cultivation experiments because of suboptimal light intensity and the presence of sucrose in the substrate which is sufficient to support the biomass increase. Since HO treated plants were greener than the control plants, it is however possible that the photosynthetic apparatus was improved upon the treatment. The photosynthesis intermediates aspartate and malate ^37,38^ present in Belgian endive water extract may help in boosting primary metabolism and thereby promote plant growth. Aspartate has been shown to improve nitrogen use efficiency in potatoes with a low supply of nitrogen ^39^. An increase in nutrient uptake, and in particular nitrogen fertilizer may stimulate the accumulation of photosynthesis proteins and contribute to plant growth. Gelatin hydrolysate, for instance, has been reported to provide a sustained source of nitrogen for cucumber growth ^40^. This biostimulant increased the expression of genes encoding for amino acid permeases (AAP3, AAP6) as well as of amino acids and nitrogen transporters ^6,40^. Increased nitrogen uptake is also associated with cell expansion that leads to elongation of the stems and petioles, effects that were also observed when plants were treated with HO.

The effect of Belgian endive root extract on root growth was more complex in that stimulating and inhibiting activity was associated with the different fractions. In addition, primary root growth is directly correlated with lateral root formation while it is inversely correlated with adventitious rooting ^41^. While HO stimulated primary root elongation (and hence also lateral root formation) at the lowest concentration, the ethanol extract (EH) strongly inhibited elongation. The growth and development of the primary and lateral roots are strongly controlled by the environment and exogenous signals including the presence of phosphate and nitrate mineral fertilizers are determining factors ^42^. Adventitious root formation is also controlled by environmental factors, yet these can be of a very different nature, such as wounding, and its regulation is likely very different from lateral root initiation and primary root growth ^43^. Despite differences in regulatory signaling pathways, the hormone auxin plays a central role in the different root structures ^44,45^. This is for instance illustrated by key auxin regulators such as *monopteros,* mutants of which were shown to lack embryonic primary root development, yet were shown to display normal adventitious root formation ^46^. Auxin is produced in the aerial part of the plant in the shoot apex and is transported basipetally to control root architecture ^47–50^. Alterations in auxin transport capacity, therefore, will have a strong impact on root growth. Since HO treatments had the opposite effect, auxin transport may have been stimulated, or alternatively, it may have enhanced local auxin synthesis ^51^.

Further characterization of the HO extract and subfractions will be critical before embarking on more detailed studies of the mode of action. Our initial efforts to purify the compounds causing bioactivity based on water-solvent partitioning were only partially successful. The liquid-liquid partitioning yielded four organic fractions (F1 – F4) and four aqueous fractions (F5 – F8). A principal component (PCA) analysis of the root and shoot effects of these fractions revealed that the aqueous fractions were much more enriched for root and shoot stimulating compounds compared to the organic fractions. Aqueous fractions are proven to be effective in the recovery of bioactive compounds during extractions ^52^. Likewise, water extracts enhance yield and improve root and shoot growth ^17^. For instance, water extracts of borage plants enhanced the yield, leaf pigment, and phenolic content of lettuce ^36^. Similarly, compost organic matter dissolved in water exhibited potential bioactivity with increased root and shoot growth, and increased enzymatic activity of nitrogen metabolism in maize ^17,53^. Clearly, the bioactivity of water-derived extracts is effective, consistent with our results.

To identify compounds putatively involved in bioactivity, the complex water fractions, F5-8 were analyzed using NMR spectroscopy which is highly suitable for characterization of complex water extracts from plants ^54–57^. Analysis of ^1^H and ^13^C spectra identified primary carbohydrates, malate, and choline, in all aqueous fractions. Since all aqueous fractions were active, these molecules are candidate bioactive ingredients. Choline has been shown to increase the rate of photosynthesis in wheat protoplasts ^58^. The salt, choline chloride, is reported to display plant growth regulatory activity in combination with chlorocholine chloride (CCC or chlormequat), an inhibitor of cyclases copalyl-diphosphate synthase and ent-kaurene synthase involved in the early steps of gibberellin biosynthesis ^59,60^). Primary carbohydrates and malate which were also found in all fractions, are not acting as growth regulators and therefore are less likely to show activity when applied in diluted concentration.

Malate is one of the organic acids considered (including citrate) a plant growth promoter, that can be utilized for low light cultivation ^61^. Malic acid is involved in several functions in plants, including respiration, nutrition, stomatal aperture, and growth ^62–64^. Reports showed that malate promotes plant growth and photosynthesis capacity - by increasing photosynthetic pigments - under normal and metal stress conditions. Darandeh and Hadavi reported in 2012 ^65^ that malate, when foliar sprayed to *Lilium* cv. Brunello, significantly increased their chlorophyl content. Likewise, malate increased the chlorophyll content of *Salix variegata* under cadmium stress. The increment was hypothetically related to the regulation of genes encoding enzymes responsible for pigment synthesis and decomposition ^38,66^. The ability of malate to alleviate metal stress has been reported in other plants including *Miscanthus saccharflorus* ^62^, *Zea mays* ^67^, sunflower ^68^, alfalfa, and white lupin ^64^. It will be interesting to test these aqueous fractions under (metal) stress conditions.

## 5. Conclusions

Belgian endive root extract contains hydrophilic and hydrophobic compounds that stimulate root and shoot growth in two unrelated species. The consistency of the bioactivity over two separate harvest years encourages further studies to identify the active compounds and determine the mode of action. In addition, it will be of interest to examine whether the compounds are active in field conditions. This will reassure a higher value of Belgian endive as a source for biostimulant production.

## 6. Patents

A European Patent Application EP21167916.2 has been filed.

## Acknowledgments

We wish to thank Versalof (http://www.versalof.be) for supplying Belgian Endive Forced Roots; and Dr. Maaike Perneel (CropFit, Ghent University), Mrs. Patricia Delaere, Mr. Christophe Petit, and Mrs. Ellen van Gysegem (Horticell Lab, Ghent University) for their technical and administrative support.

## Author Contributions

Conceptualization, D.G., B.V.D., S.M., and S.W.; validation, D.G., B.V.D., S.W., and A.R.; formal analysis, H.Y.O., P.M., and A.R.; investigation, H.Y.O., J.L., L.X., H.T.K., N.B., P.M., and N.M.; resources, D.G., B.V.D., S.M., and S.W.; data curation, A.R.; writing—original draft preparation, H.Y.O.; writing—review and editing, D.G., and A.R.; visualization, H.Y.O.; supervision, D.G., B.V.D., S.M., and S.W.; project administration, A.R.; funding acquisition, D.G., B.V.D., S.M., and S.W. All authors have read and agreed to the published version of the manuscript.

## Additional information

### Supplementary materials

The following are available in the supplementary file. Figure S1: Schematic representation of the sequential extraction procedure used in the preparation of the four Belgian endive forced roots extracts. Table S1: Technical details of the fractionation procedure of the 2018 Belgian endive forced roots water (HO) extract. Figure S2: *Arabidopsis* root architecture stimulation upon treatment with 2018 crude endive extracts (EH, EA, HE). Figure S3: Pictures showing the growth of *Arabidopsis* root and shoot stimulation 10 days after treatment with 2020 crude “HO” endive extract. Figure S4: Graphs showing the consistent positive effect of low, mid, and high doses of HO on *Arabidopsis* plants from two different production and extraction periods (2018 and 2020). Figure S5: Graphical representations showing the fractions of HO on *Plectranthus* explants. Figure S6: Principal component analysis (PCA) of *Arabidopsis* phenotypes under organic or aqueous fractions treatments. Figure S7: ^1^H NMR (D_2_0, ref: DSS. pH 7, 400 MHz) spectrum of F5 (a) and F6 (b) with representative peaks. Figure S8: ^13^C NMR (D_2_0, ref: DSS. pH 7, 100.6 MHz) spectra of F5, F6 (a) and F7, F8 (b) in the 0 – 110 ppm region. Table S2: ^13^C NMR chemical shift assignment of identified metabolites and their structure.

