## supplementary file for "Belgian endive-derived biostimulants promote shoot and root growth *in vitro*"

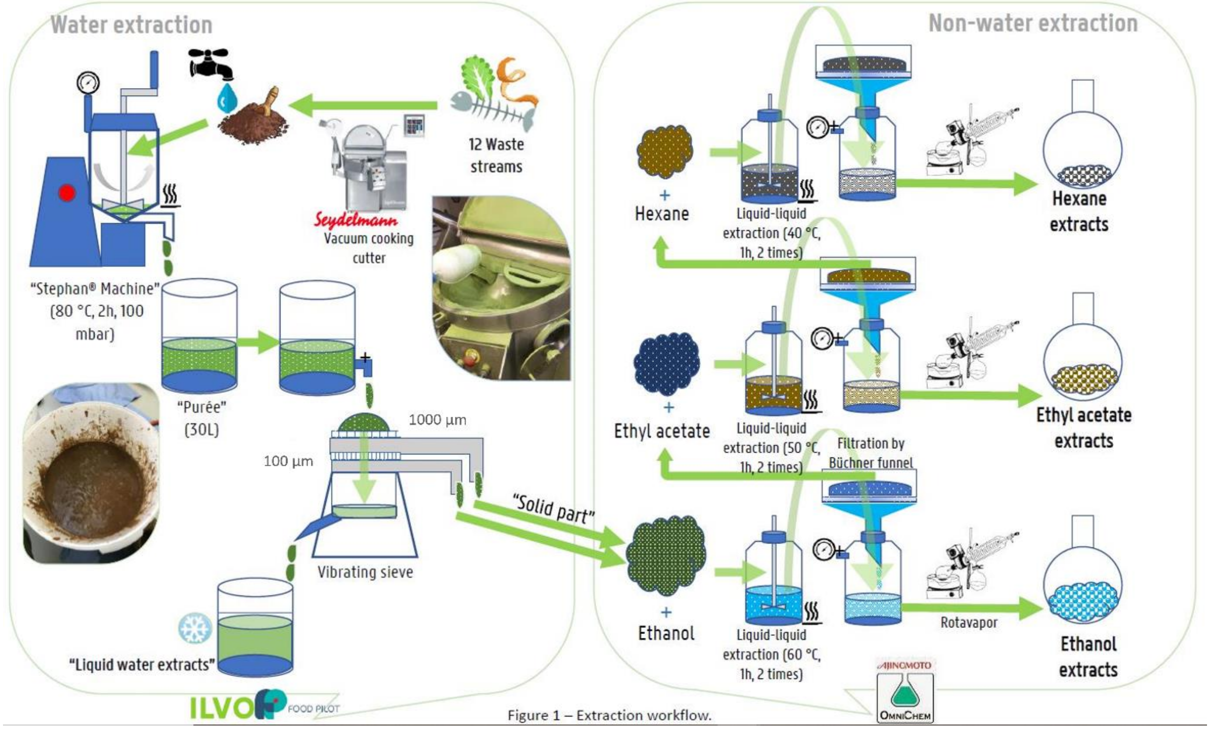


**Figure S1**. Schematic representation of the sequential extraction procedure used in the preparation of the four Belgian endive forced roots extracts. Liquid water (HO) extract, dried ethanol (EH) extract, dried ethyl acetate (EA) extract, and dried hexane (HE) extract.

Table S1. Technical details of the fractionation procedure of the 2018 Belgian endive forced roots water (HO) extract.

| **Extract volume** before **concentration (mL)/pH** | **Extract volume** after **concentration (mL)/pH** | **Extract aliquots (mL)** | **pH** | **$H_2O$ (mL)** | **Ethyl acetate (mL)** | | | **Toluene (mL)** | | | **Fraction yield (mg[1] or mL[2])** | **Fraction name** |
|---|---|---|---|---|---|---|---|---|---|---|---|---|
| 11000/6.25 | 2105/6.02 | 500 | 3 | 400 | 750 | 700 | 750 | | | | 1026[1] | F1 |
| | | 575 | 10 | | 750 | 700 | 750 | | | | 506[1] | F2 |
| | | 490 | 3 | | | | | 750 | 700 | 700 | 397[1] | F3 |
| | | 540 | 10 | | | | | 750 | 750 | 700 | 100[1] | F4 |
| | | | | | | | | | | | 2625[2] | F5 |
| | | | | | | | | | | | 2800[2] | F6 |
| | | | | | | | | | | | 2560[2] | F7 |
| | | | | | | | | | | | 2785[2] | F8 |

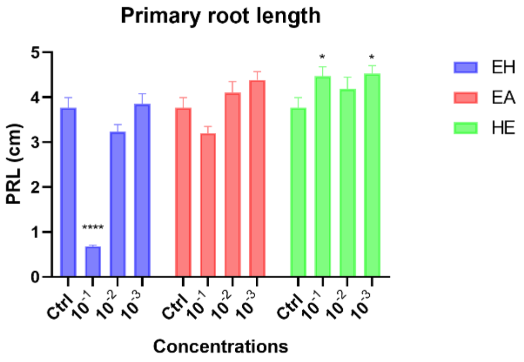


(a)

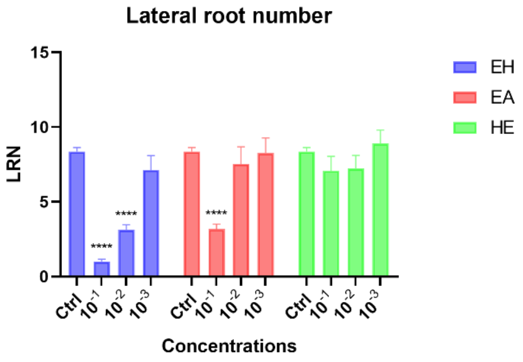


(b)

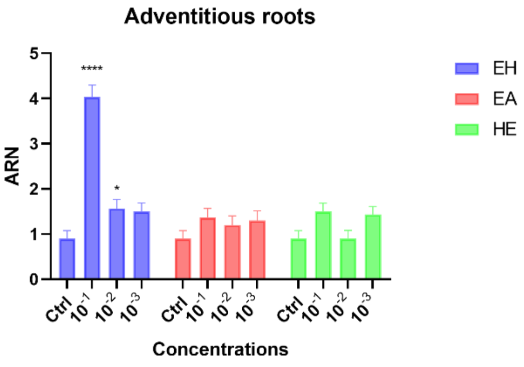


(c)

**Figure S2**. *Arabidopsis* root architecture stimulation upon treatment with 2018 crude endive extracts (EH, EA, HE). Graphical representations showing the effect of 10-, 100-, and 1000-times dilution of EH, EA, and HE on *Arabidopsis* plants. (a) showed the effect on primary root length; (b) showed the effect on lateral root number; and (c) showed the effect on adventitious roots. PRL: primary root length, LRN: lateral root number; ARN: adventitious root number. Data represent the average of three biological and ten technical replicates per bar (30 seedlings in total, 10 per replicate). Error bars represent standard mean error (SEM). Asterisks indicate significant differences (Dunnett's multiple comparison test. * $p<0.05$, ** $p<0.01$, *** $p<0.001$, **** $p<0.0001$).

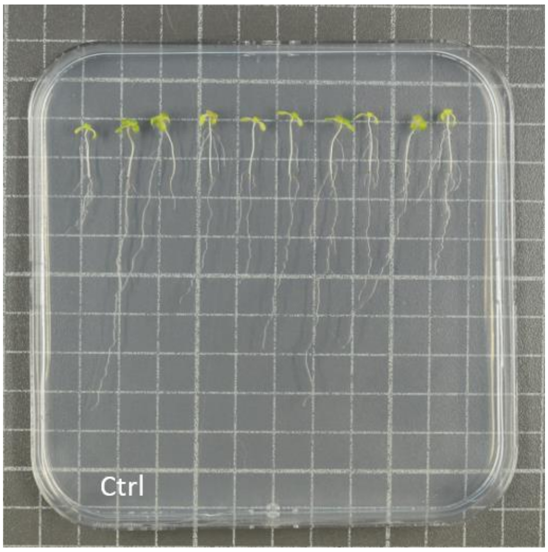


(a)

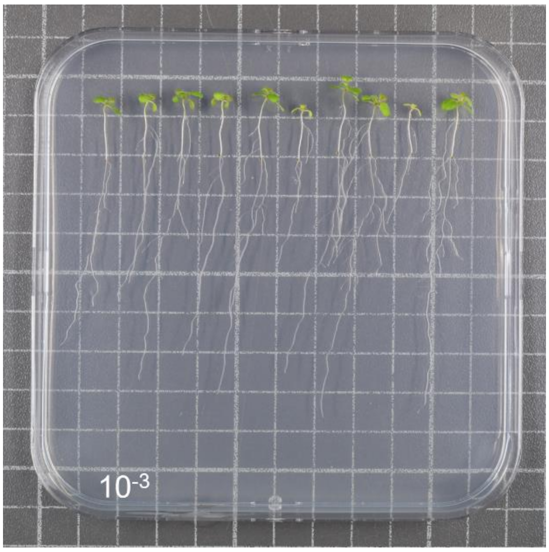


(b)

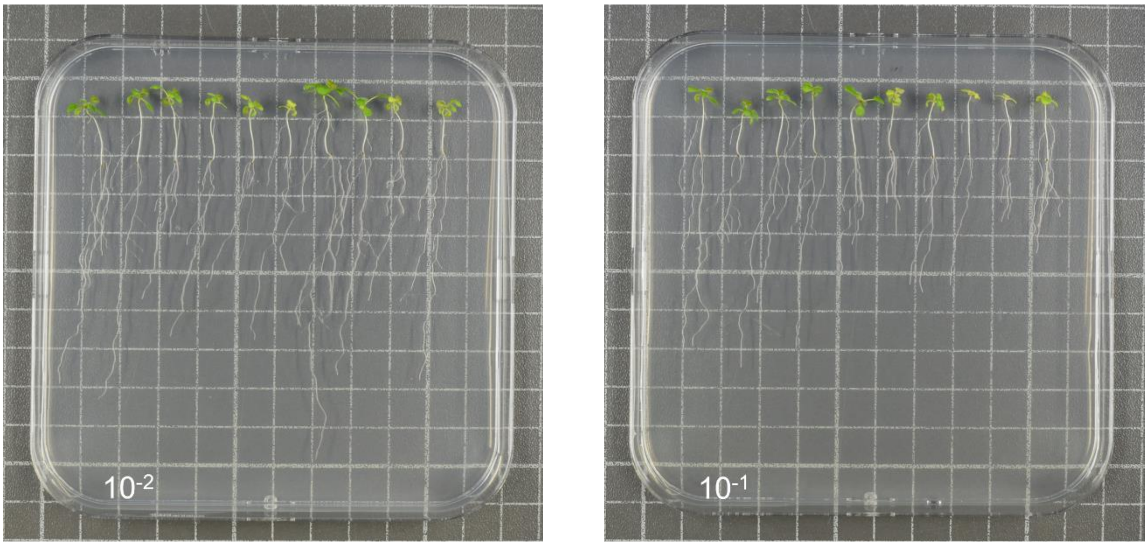


(c) (d)

**Figure S3**. Pictures showing the growth of *Arabidopsis* root and shoot stimulation 10 days after treatment with 2020 crude “HO” endive extract. The pictures show the morphology of (a) control (Ctrl), (b) low, (c) mid, and (d) high doses of HO treated plants.

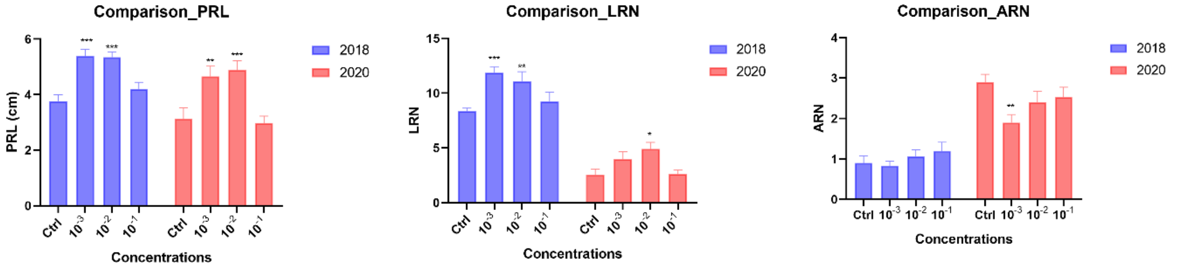


(a) (b) (c)

**Figure S4**. Graphs showing the consistent positive effect of low, mid, and high doses of HO on *Arabidopsis* plants from two different production and extraction periods (2018 and 2020). (a) the effect on primary root length (PRL); (b) the effect on lateral root number (LRN); and (c) the effect on adventitious root number (ARN) of Arabidopsis compared to the control (Ctrl) plants. Data represent the average of three biological and ten technical replicates per bar (30 seedlings in total, 10 per replicate). Error bars represent standard mean error (SEM). Asterisks indicate significant differences ($p<0,05$) between negative control (Ctrl) and treatments ($10^{-3}$, $10^{-2}$, and $10^{-1}$ of HO) in respective groups (experiments year) according to Tukey’s multiple comparison test * $p<0.05$, ** $p<0.01$, *** $p<0.001$, **** $p<0.0001$.

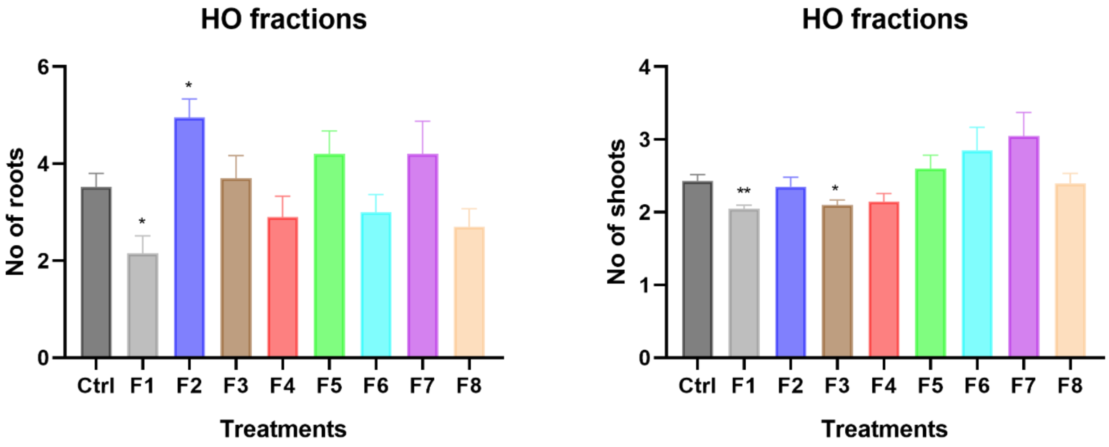


(a) (b)

**Figure S5.** Graphical representations showing the fractions of HO on *Plectranthus* explants. (a) shows the effect on the root numbers and (b) shows the effect on the shoot numbers. Data represent the average of two biological and five technical replicates per bar. Error bars represent standard mean error (SEM). Asterisks indicate significant differences (Dunnett's multiple comparison test. * p<0.05, ** p<0.01, *** p<0.001, **** p<0.0001).

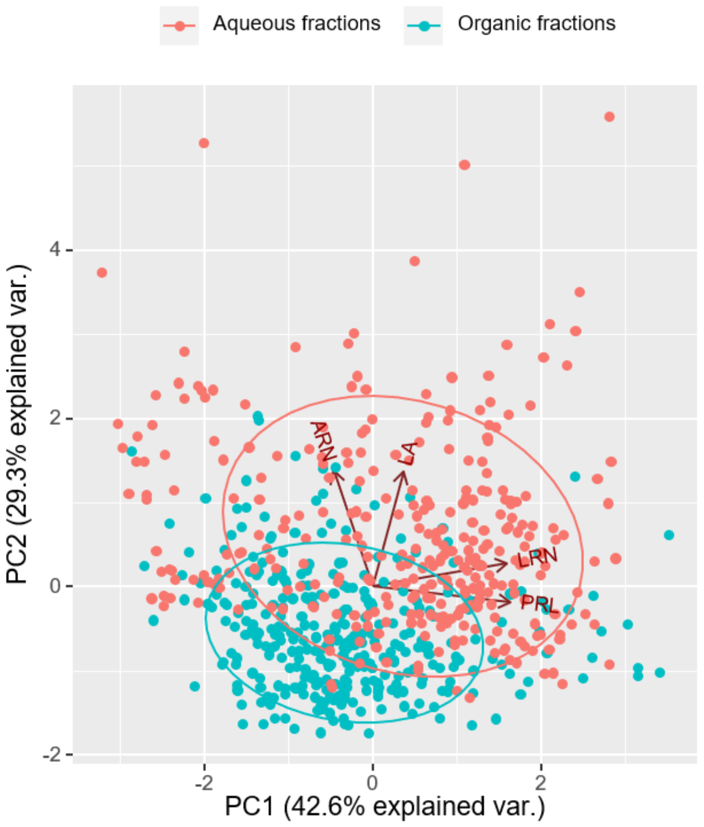


**Figure S6.** Principal component analysis (PCA) of *Arabidopsis* phenotypes under organic or aqueous fractions treatments. The graph shows the PCA biplot of individual samples to PC 1 and PC 2. Colors indicate fractions treatment (blue: organic fractions and red: aqueous fractions). The plant phenotypes are adventitious root number (ARN), leaf area (LHA), lateral root number (LRN), and primary root length (PRL).

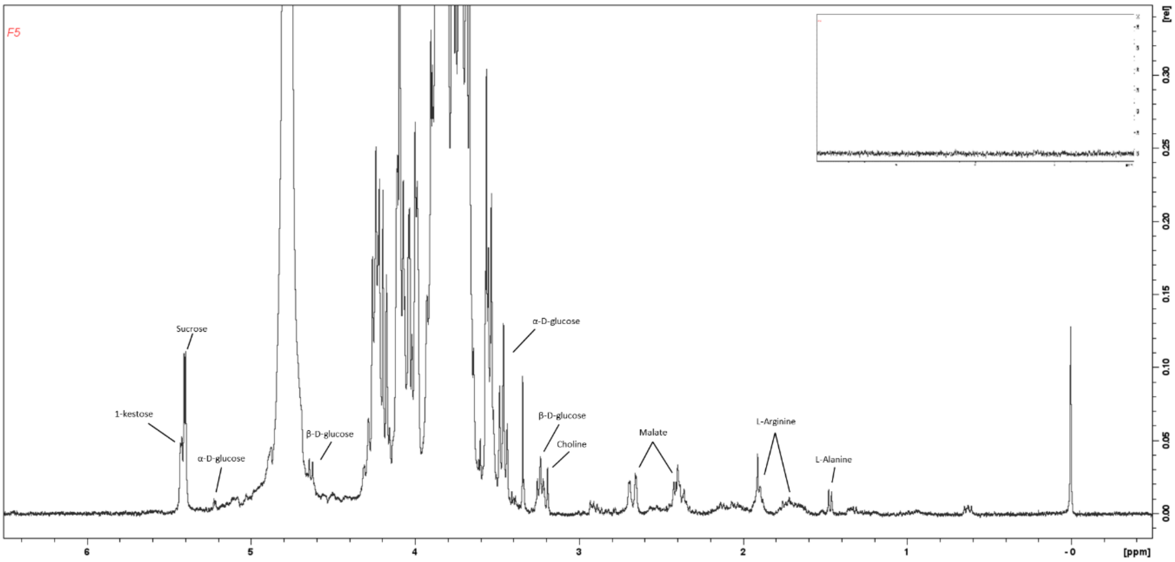


(a)

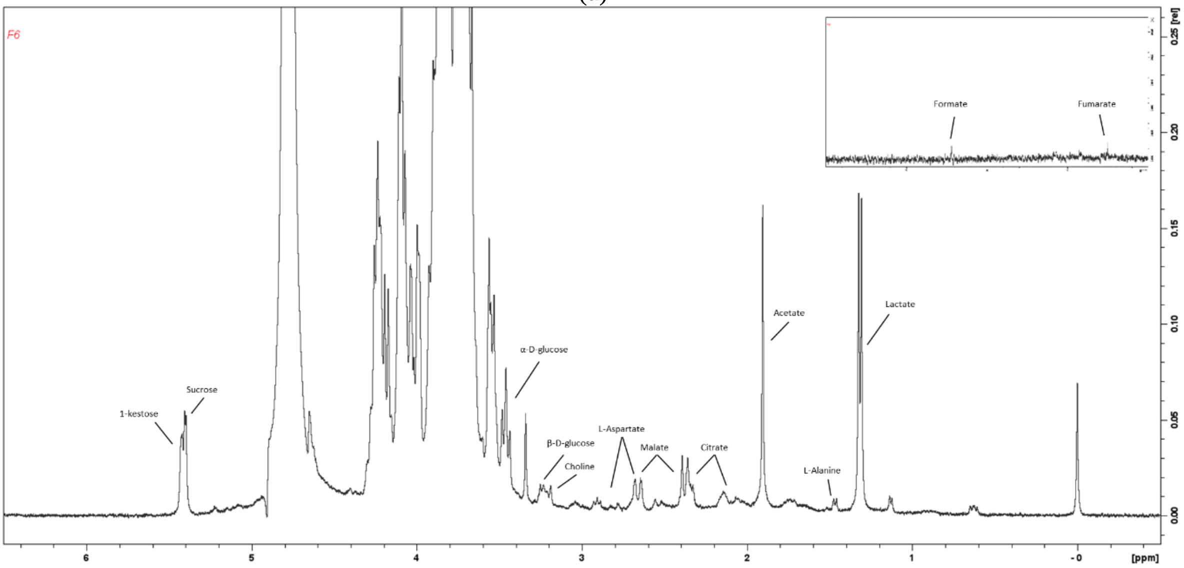


(b)

**Figure S7.** [1]H NMR ($D_20$, ref: DSS. pH 7, 400 MHz) spectrum of F5 (a) and F6 (b) with representative peaks. On top right, zoom of region 6 – 10 ppm.

(a)

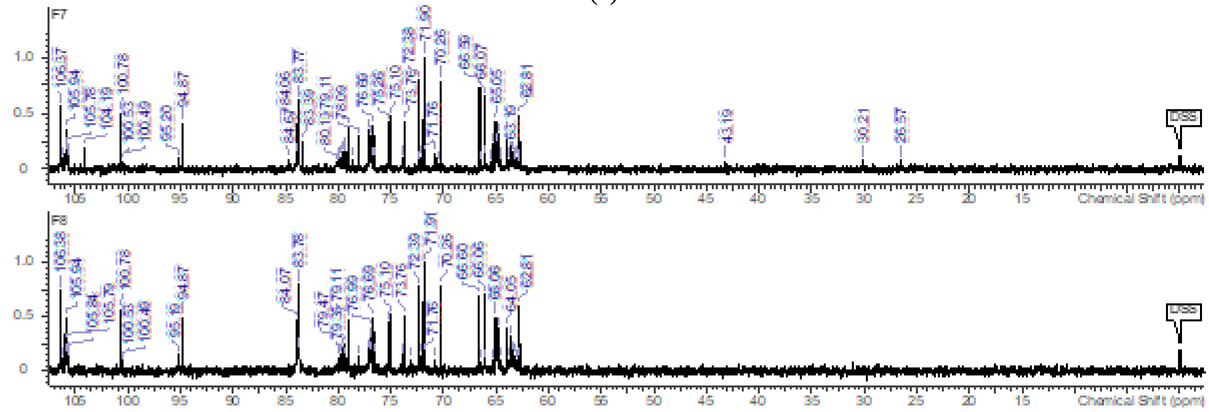


(b)

**Figure S8.** $^{13}$C NMR ($D_20$, ref: DSS. pH 7, 100.6 MHz) spectra of F5, F6 (a) and F7, F8 (b) in the 0 – 110 ppm region.

Table S2. $^{13}C$ NMR chemical shift assignment of identified metabolites and their structure.

| Metabolites | Residue | Carbon (number) | Chemical shift (ppm) | Structure |
|---|---|---|---|---|
| **Sucrose** | | 1 | 106.4 | 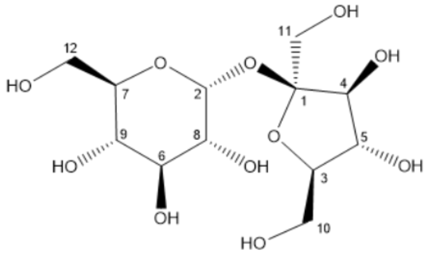 |
| | | 2 | 94.9 | |
| | | 3 | 84.0 | |
| | | 4 | 79.1 | |
| | | 5 | 76.7 | |
| | | 6 | 75.1 | |
| | | 7 | 75.1 | |
| | | 8 | 73.8 | |
| | | 9 | 71.9 | |
| | | 10 | 65.1 | |
| | | 11 | 64.0 | |
| | | 12 | 62.8 | |
| **Fructofuranose (α and β)** | | 1 | 104.2 | 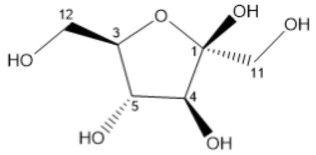 |
| | | 2 | 100.8 | |
| | | 3 | 83.4 | |
| | | 4 | 78.1 | |
| | | 5 | 77.1 | |
| | | 6 | 72.4 | |
| | | 7 | 71.9 | 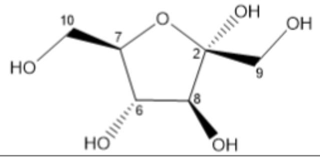 |
| | | 8 | 70.3 | |
| | | 9 | 66.6 | |
| | | 10 | 66.1 | |
| | | 11 | 65.4 | |
| | | 12 | 65.1 | |
| **1-kestose** | $F_1$ | 1 | 63.6 | 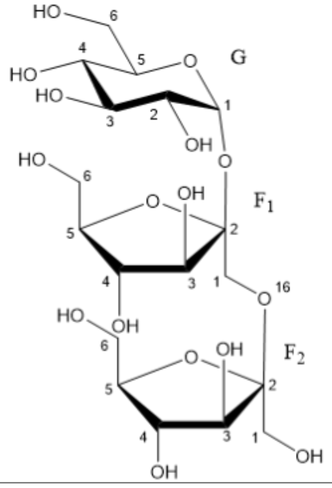 |
| | $F_1$ | 2 | 105.9 | |
| | $F_1$ | 3 | 79.5 | |
| | $F_1$ | 4 | 76.6 | |
| | $F_1$ | 5 | 83.9 | |
| | $F_1$ | 6 | 64.9 | |
| | $F_2$ | 1 | 63.1 | |
| | $F_2$ | 2 | 106.4 | |
| | $F_2$ | 3 | 79.5 | |
| | $F_2$ | 4 | 77.1 | |
| | $F_2$ | 5 | 83.8 | |
| | $F_2$ | 6 | 65.1 | |
| | G | 1 | 95.2 | |
| | G | 2 | 73.9 | |
| | G | 3 | 75.3 | |
| | G | 4 | 71.9 | |
| | G | 5 | 75.1 | |
| | G | 6 | 62.8 | |
| **L-arginine** | | $\gamma - CH_2$ | 26.6 | 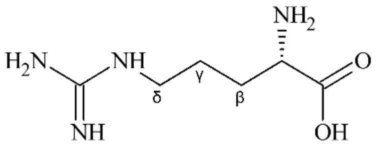 |
| | | $\beta - CH_2$ | 30.3 | |
| | | $\delta - CH_2$ | 43.3 | |
| **Lactate** | | $CH_3$ | 22.8 | 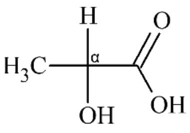 |
| | | α-CH | 71.2 | |
| | | -COOH | 185.3 | |
